# Spatiotemporal dynamics across visual cortical laminae support a predictive coding framework for interpreting mismatch responses

**DOI:** 10.1101/2023.04.17.537173

**Authors:** Connor G. Gallimore, David Ricci, Jordan P. Hamm

## Abstract

Context modulates neocortical processing of sensory data. Unexpected visual stimuli elicit large responses in primary visual cortex (V1) -- a phenomenon known as deviance detection (DD) at the neural level, or “mismatch negativity” (MMN) when measured with EEG. It remains unclear how visual DD/MMN signals emerge across cortical layers, in temporal relation to the onset of deviant stimuli, and with respect to brain oscillations. Here we employed a visual “oddball” sequence – a classic paradigm for studying aberrant DD/MMN in neuropsychiatric populations – and recorded local field potentials in V1 of awake mice with 16-channel multielectrode arrays. Multiunit activity and current source density profiles showed that while basic adaptation to redundant stimuli was present early (50ms) in layer 4 responses, DD emerged later (150-230ms) in supragranular layers (L2/3). This DD signal coincided with increased delta/theta (2-7Hz) and high-gamma (70-80Hz) oscillations in L2/3 and decreased beta oscillations (26-36hz) in L1. These results clarify the neocortical dynamics elicited during an oddball paradigm at a microcircuit level. They are consistent with a predictive coding framework, which posits that predictive suppression is present in cortical feed-back circuits, which synapse in L1, while “prediction errors” engage cortical feed-forward processing streams, which emanate from L2/3.

## Introduction

The neocortex processes incident sensory information not in isolation, but in the rich spatiotemporal context in which it presents. In a predictive coding framework, internal models of the environment, encoded within high-level brain regions, serve to modulate sensory cortical processing, suppressing low-level responses to expected or redundant sensory data (Friston 2005, 2018; Bastos et al. 2012). On the other hand, unexpected or novel stimuli evoke large sensory cortical responses. Such augmented responses are thought to reflect perceptual “prediction-errors” in cortical circuits, signaling a mismatch between experienced vs expected sensory information (Näätänen 1990; Jordan and Keller 2020; Hamm et al. 2021). Prediction-errors are fed forward, from lower to higher brain regions, to update internal models of the environment (Friston 2005), thus forming a critical component in perception, learning, and cognition (Friston 2018).

Major psychotic disorders like schizophrenia involve persistent perceptual symptoms such as hallucinations and aberrations, but also lower-level abberations in sensory cortical processing which, though subtle, are highly reliable across patients and time, emerge early in the disease process, and are heritable (Javitt 2009; Javitt and Freedman 2015). The predictive coding theory of cortical responses (Sterzer et al. 2018) provides a potent framework for understanding these symptoms and sensory processing abnormalities in terms of concrete and quantifiable aberrations in neural systems (e.g. aplastic neural circuits due to weak prediction-errors generated in lower-level neural circuits). Further, evidence of aberrant predictive processing is well documented in schizophrenia (SZ). The mismatch negativity (MMN) is an electroencephalographical (EEG) scalp potential experimentally elicited during a classic “oddball” paradigm, which involves a train of “redundant” or “standard” stimuli (e.g. a repeated tone or bar orientation) with rare “deviant” or “target” stimuli interspersed (e.g. a different tone or orientation). Specifically, the MMN is a scalp potential elicited 150-200ms after the onset of a “deviant” stimulus. Individuals with SZ exhibit reductions in MMN in the auditory domain (Light and Näätänen 2013; Näätänen et al. 2015) as well as the visual domain (Farkas et al. 2015; Kremláček et al. 2016). Though MMN primarily reflects an early sensory cortex signal (Garrido et al. 2009) elicited in the absence of direct attention or task demands (Light and Näätänen 2013), deficits in MMN correlate strongly with higher-level functional (Light and Braff 2005) and cognitive difficulties in the disease (Revheim et al. 2014), and predict conversion to psychosis in at-risk individuals (Shin et al. 2009; Perez et al. 2014). MMN deficits are thus thought to index aberrations in elementary predictive processing (Sterzer et al. 2018) which undermine how individuals perceive and cognitively relate to the world (Javitt and Freedman 2015).

Almost three decades of work highlight MMN as one of the best replicated biomarkers of sensory processing dysfunction in SZ (Erickson et al. 2016; Kremláček et al. 2016), yet the brain dynamics underlying its generation are only partially understood, limiting the clinical utility of this otherwise robust biomarker. Further, whether MMN and other cortical dynamics present during a passive sensory oddball paradigm are consistent with the “predictive coding” framework has been proposed (Friston 2005; Hamm et al. 2021), but direct evidence is incomplete. Some recent work in rodents suggests that neurons in sensory cortices exhibit genuine “deviance detection” (DD) responses to deviant stimuli during the oddball paradigm (Hamm and Yuste 2016; Harms et al. 2016). Such DD Reponses share paradigmatic and temporal features analogous to the MMN and are thusly considered a cell-level or intracortical analogue of MMN measurable in rodents and non-human primates (Ross and Hamm 2020). DD has been proposed to represent a sensory prediction error (Friston 2005). Deviance detecting neurons in V1 are mostly present in superficial areas (Hamm et al. 2021), which are known to project primarily cortico-cortically (often to higher brain areas), consistent with “prediction-error” propagation in the predictive coding framework (Bastos et al. 2012). However, whether oddball-evoked spatiotemporal dynamics across other layers are consistent with a predictive coding framework – reflecting top-down predictions and integrative prediction-errors with distinct spatial and time-frequency signitures– has not been established.

While human-like MMN scalp potentials are difficult to directly equate to EEG or local field potentials (LFPs) in rodents, both human MMN and rodent cortical DD responses to auditory oddballs exhibit local field potential energy in the low theta frequency band (2-8 Hz) (Javitt et al. 2018; Lee et al. 2018). This is consistent with the notion that MMN indexes a feed-forward prediction-error signal (Bastos et al. 2012), as oscillations in this frequency domain have been demonstrated to index feed-forward connections in cortical networks. It remains unclear whether this pattern holds true for *visual* MMN and oddball-elicited activity as well, a potentially valuable insight given that SZ involves macro and microanatomical pathology in visual cortices (Hashimoto et al. 2008; Türközer et al. 2022), with visual MMN reduced in SZ (Kremláček et al. 2016).

Visual MMN-like signals have been recently studied at the cellular level in mouse models (Hamm and Yuste 2016; Hamm et al. 2021), showing genuine “deviance detection” characteristics necessary for translational work (Harms et al. 2016). Further, beta-band oscillations in cortical systems reflect feed-back “predictions” (Bastos et al. 2015). The role of beta-band synchronizations, as well as dynamics in other frequency bands, in sensory cortical MMN or DD has not been established, but could provide further evidence for MMN as a predictive coding biomarker.

Thus, the goals of this study were i) to provide a deeper understanding of the cortical electrophysiological dynamics present during a visual oddball paradigm at the level of a cortical column, highlighting the laminar and neurooscillatory signitures of DD and other predictive processes (e.g. stimulus specific adaptation). We expected V1 DD to correspond to increased theta-band power (as has been identified in rodent auditory cortical DD) and increased neural spiking in layer 2/3 neural activity, and we expected SSA to correspond with decreases in early evoked neural activity in granular layers. Whether and how other oscillatory frequency bands and layers are modulated by deviance and redundancy was a more open question. Therefore, sought ii) to test whether neural firing, synaptic currents, and oscillatory dynamics across layers within a cortical column are consistent with a predictive coding framework with regard to feed-forward and feed-back circuity (Bastos et al. 2012, 2020). We recorded extracellular LFPs across 750µm of depth in primary visual cortex (V1) with a 16-channel multielectrode shank, analyzing how processing of the same stimulus across different contexts (i.e the oddball and control sequences) is reflected in multiunit activity (an index of aggregate neural spiking) and neural oscillatory synchrony within and across granular, supragranular, and infragranular laminae. We focused on DD: enhanced brain responses to a given stimulus when it is the “oddball” (rare and contextually deviant) relative to when it is present in a “many-standards” control sequence (rare, but not contextually deviant).

## Materials and Methods

### Animals, Surgery, and Training

All procedures were conducted in accordance with Georgia State University IACUC guidelines. Adult male and female mice (10-24 weeks; 7M, 3F) with VIP-cre, SST-cre, or Vglut-cre genetic backgrounds (Jax# 010908 (Taniguchi et al. 2011), 013044 (Taniguchi et al. 2011), 023527 (Harris et al. 2014), respectively) were used for LFP experiments. All mice were implanted with custom titanium headplates (permitting head-fixation in front of a stimulus monitor) and a titanium reference screw overlying the cerebellum at least one week before experiments. After recovery, mice underwent 2-3 consecutive days of head-fixation training prior to experiments. During training, mice were exposed to sequences of visual moving grating stimuli (described below) for acclimation. On the day of experiments, a small craniotomy above the left V1 was performed for acute electrode insertion.

### Visual Stimuli

As previously described (Hamm et al. 2021), full-field square-wave gratings (100% contrast; .08 cycles per degree) of 30, 45, 60, 90, 120, 135, 150, and 180-deg orientations were created with MATLAB Psychophysics Toolbox (Brainard 1997; Pelli 1997) and displayed on an LCD monitor positioned ≈20cm from the right eye (500ms duration; ISI of ∼500ms black screen). Oddball experiments consisted of two sequential runs: control and test. For the “many-standards” control sequence, all 8 orientations were presented randomly with approximately equal likelihood (≈12.5% probability). Afterwards, two stimuli separated by 90-deg (e.g. 45-deg and 135-deg) were selected for redundant (≈87.5% probability) and deviant (≈12.5%) presentation in the oddball test sequence, and then “flip-flopped” within the same run, such that the previously redundant stimulus served as the deviant and vice versa. This procedure ensured at least two stimuli participated in all three sensory contexts: equiprobable (control; C), high likelihood (redundant; R), and rare (deviant; D). Sequences lasted 5-6 minutes each, with ≈250 trials in the control run and ≈150 in each test run (Figure 1a-b).

**Figure 1.**
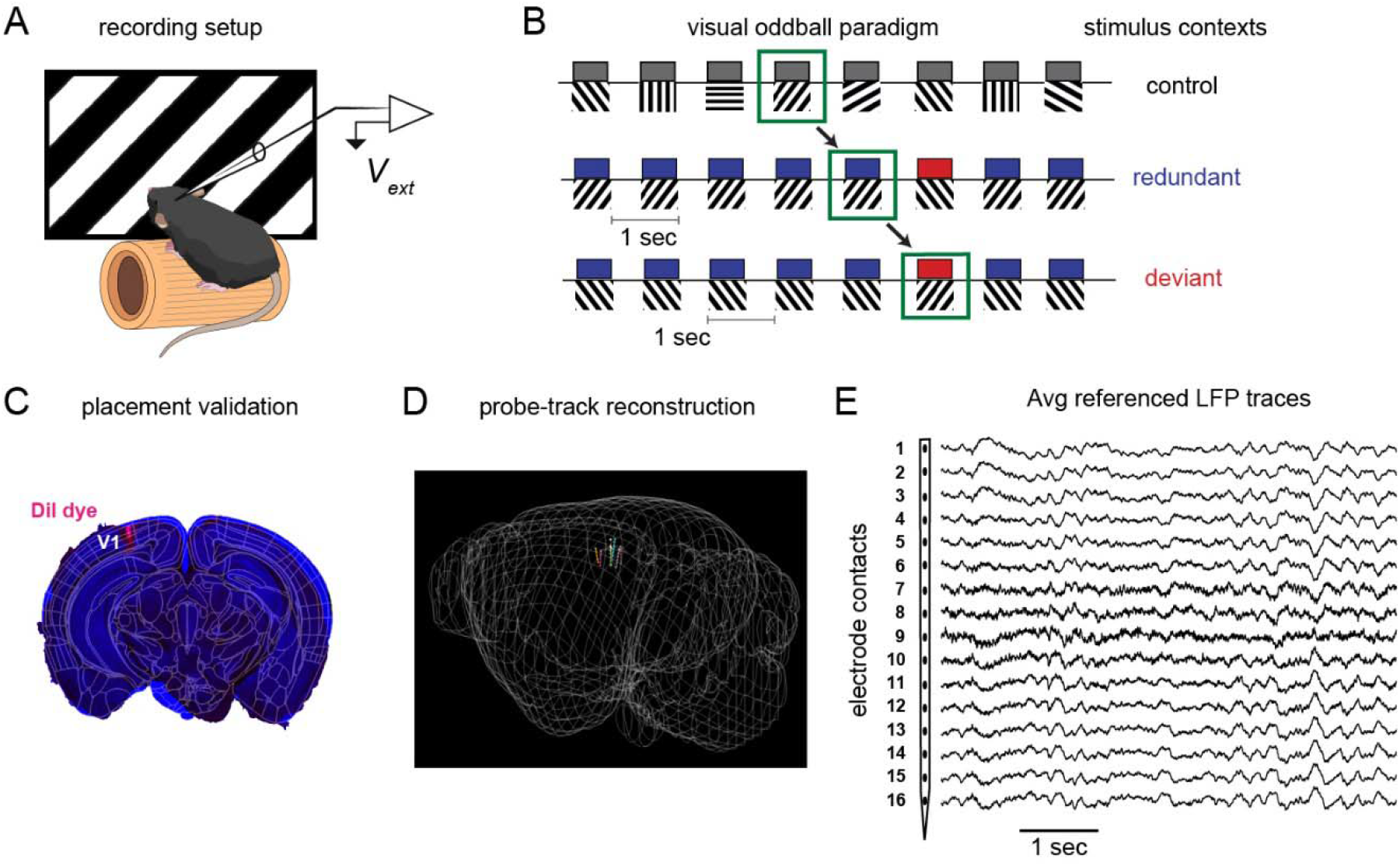
Overview of visual oddball paradigm and data collection methods. **a)** Depiction of awake mouse on treadmill with an extracellular recording probe in V1. b) Progression of sensory oddball paradigm with stimulus probabilities to the right of their respective context. c) estimated trajectories from dye tracks in histological sections registered to the Allen CCF; 7 of 10 electrode positions reconstructed in d). e) Mean-subtracted raw LFP traces for the 16 electrode contexts spanning 750um ventral V1 surface.

### Electrophysiology: collection and processing

Intracortical electrical signals were recorded from a custom designed 16-channel NeuroNexus probe (750μm length, 50μm inter-contact distance; A1×16–3mm50–177; Ann Arbor, MI) inserted perpendicularly into left V1 at 100μm/min until the dorsal-most electrode was just below the dura (deduced from real-time signals). Prior to insertion, probes were submerged in Dil dye for post-hoc anatomical validation (figure 1c-d). Signals were digitized at 10kHz and processed as either local field potentials (LFP) or multiunit activity (MUA). All analyses were performed using MATLAB (The Math Works 2020). For LFP analyses, we low-pass filtered (<110Hz), and then either a) converted to current source densities as previously described (Hamm and Yuste 2016; Hamm et al. 2020) (CSDs; 2^nd^ spatial derivative of voltage, gaussian-smoothed with a 5-point Hamming window) or b) converted to the time-frequency domain for analysis of single trial induced power or inter-electrode phase synchrony. For analysis of MUA, we processed data in line with past work (Kirchberger et al. 2021). Channels were filtered between 500 and 4000 Hz and common average referenced the other 15 channels to remove artifacts such as EMG activity (Ludwig et al. 2009). Activity was then rectified and smoothed in 10ms gaussian windows, and down-sampled to 1000Hz for further analysis. Consistent with i) the fact that MUA is known to reflect local neural firing activity, and ii) the fact that V1 neural responses are orientation selective, we found that MUA responses were highly variable across stimuli of different orientations/direction within mice. Thus we focused our MUA and LFP/CSD analyses on only 1 stimulus orientation per mouse: the orientation with the strongest MUA response across layers and contexts in the first 100ms after stimulus onset (i.e. 0, 90, 45, or 135 degrees). One mouse did not display clear MUA responses to any stimulus and was not included in the MUA analysis.

Multielectrode data for MUA, CSD, and time-frequency analyses were centered around putative layer 4 based on the stimulus-triggered average responses to all stimuli. Criteria for alignment to layer 4 focused on i) the contact with the earliest MUA peak and current source/sink in the CSD (Niell and Stryker 2008; Ferro et al. 2021) and ii) the more recently demonstrated shift from gamma to alpha/beta power spectra (which reflects supra-vs infra-granular layers; (Sanchez-Todo et al. 2023)). This method also correlated well with the channel at which the LFP signals flipped polarity. We then restricted analyses to the 12 electrode contacts around this point, with 7 above and 5 below (12 total). For MUA, we analysed each of the 12 contacts separately. For CSD and LFP, activity tends to be more spatially distributed and correlated, with sink/source patterns spanning multiple contacts within layers. Thus, we focused on 4 putative “layers” in concordance with anatomical literature and the Allen institute mouse brain atlas. The top 2 electrode contacts (from the top) were considered superficial/layer 1, the next 5 were considered supragranular (layer 2/3), the next 2 were considered granular (layer 4), and the last 3 were considered infragranular (mainly layer 5a). Within each “layer”, we analysed CSD as an “av-rec” montage (Javitt et al. 1996) by averaging all CSD profiles across trials within mice at the individual channel level and then taking the absolute value of all currents within a “layer” and averaging to derive a single av-rec CSD waveform for each mouse and each of the 4 layers.

For conversion to the time-frequency (t-f) domain for analysis of oscillatory power and phase-locking, we used EEGLAB (Delorme and Makeig 2004) to apply modified Morelet-wavelets to LFP data from individual trials, comprising 100 equally-spaced wavelets (2-101Hz, 1Hz resolution, linearly increasing from 0.5 cycles to 20 cycles) applied every 4ms from -250ms pre-stimulus onset to 250ms post stimulus offset. Stimulus-induced power was quantified as the across-trial average of the squared magnitudes of the absolute value of the complex output of the wavelet analysis. We subtracted a global baseline for each frequency, mouse, and layer (averaged -200-50ms pre-stimulus across all conditions and trials) in order to scale responses for easier plotting/display while avoiding contamination of baseline differences and/or variability in the inter-context comparisons. Stimulus-induced inter-electrode synchrony (IES) was calculated by quantifying the phase consistency (Hamm et al. 2020) of inter-electrode phase lags during the 50-250 (early) and 250-450ms (late) post-stimulus time period. The primary goal was to assess how stimulus-induced inter-laminar interactions relate to visual stimulus context. To adjust for indirect causes of inter-electrode phase coherence, such as shared phase coherence at two contacts with visual stimulus onset (thus, independent of the interactions between electrodes, but convolved with responses to the stimulus of interest), we calculated a “baseline” surrogate IES measure by computing coherence values between electrodes on different trials (of the same stimulus; 2 trials in the future) and subtracting that from the actual IES, resulting in a final IES measure which better reflects stimulus-related interelectrode interactions (Canolty et al. 2006). Resulting IES was summed over trials to yield a final IES spectra (dims 12x12x100) for deviant, redundant, and control contexts. IES maps (figure 4) were constructed by defining a seed channel and computing the phase-locking present between each electrode in the array from the one chosen.

### Analysis and statistics

Primary analyses focused on evoked responses to the same stimulus across the three control, redundant, and deviant contexts (figure 1b). The number of trials was held constant across conditions (between 7 and 12 across mice, depending on number of trials without artifacts).

Evaluations of MUA (12 channels) and CSD (4 layers) were carried out on spatiotemporal bins of 30ms centered around time periods of maximal average evoked responses (mostly from 50 to 260ms post-stimulus onset). Time/depth widows without strong MUA and/or CSD were not analysed (e.g. contacts 4 and 5 at 50-80ms in the MUA; Fig 2E). Our *a priori* hypotheses guided these comparisons based on the assumption that early granular activity (50-140ms) would show stimulus-specific adaptation (SSA: control greater than redundant) and later supragranular activity (141-260ms; in the range of the MMN and P300 potentials) would show deviance detection (DD; deviant greater than control). MUA analyses were one-tailed t-tests, as the direction of the hypothesized SSA and DD effects were clear (Hamm and Yuste 2016; Ross and Hamm 2020; Hamm et al. 2021). CSD analyses were two-tailed t-tests, as SSA or DD do not necessarily match up to more vs less transsynaptic activity (e.g. SSA could be associated with more inhibitory current or less excitatory, and thus a greater or lesser CSD/LFP values, depending on the layer and timepoint).

**Figure 2.**
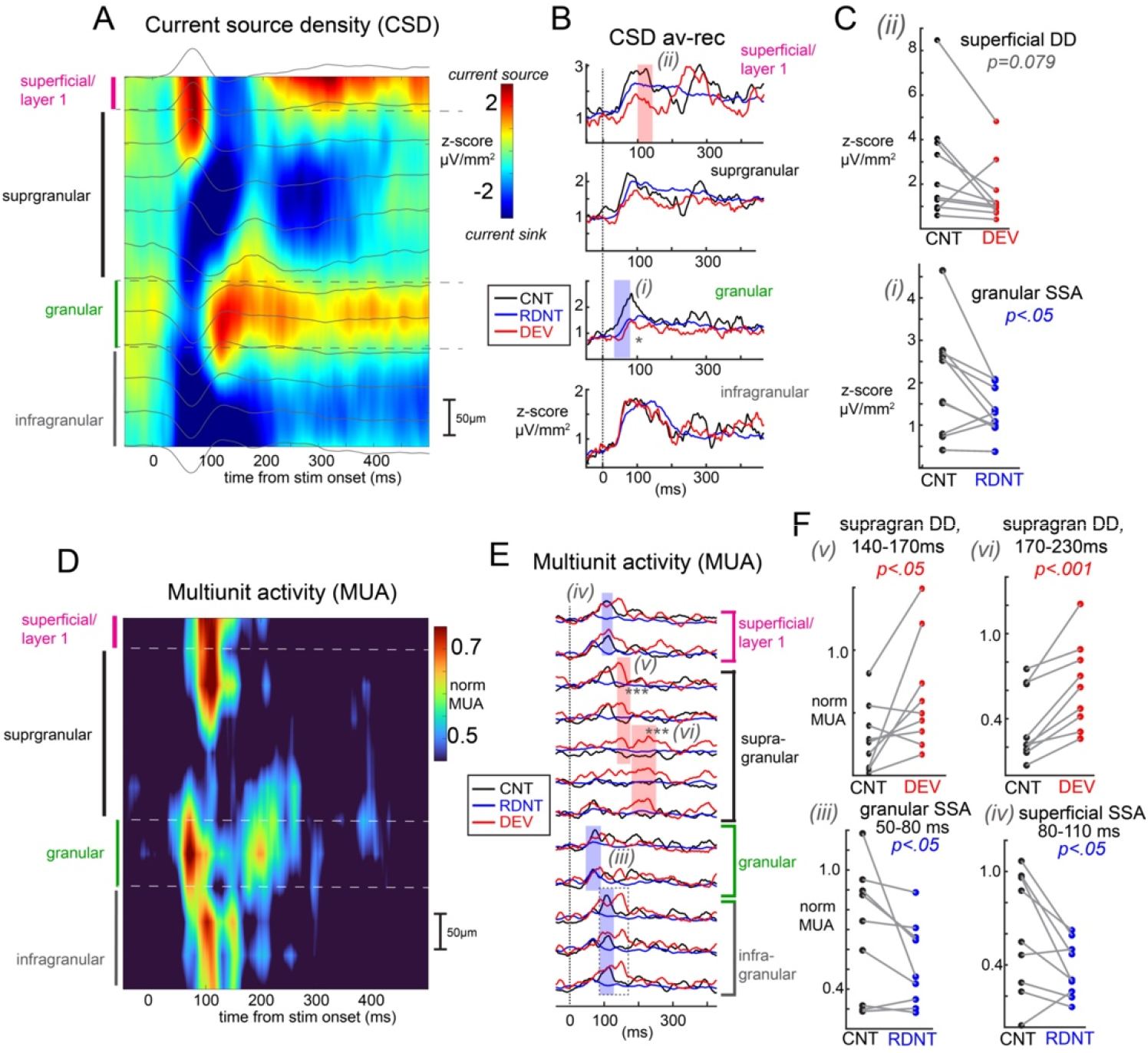
Evoked current source density and multiunit spiking segregate SSA and DD in V1. A) Current source density (CSD) spectra aligned to each recording’s putative layer 4 (granular) layer and averaged over all recordings (10 mice). Layer definitions are denoted by dotted gray lines. Background gray lines depict average LFP. B) Average within-layer rectified CSD traces, averaged across trials and 9 mice within control, redundant, and deviant contexts. Thirty millisecond time windows exhibiting significant SSA (control vs redundant) or DD (control vs deviant) are indicated with shaded rectangles. C) Scatterplots of individual recordings (mice) from shaded areas in B. D-F: same as A-C, but for multiunit spiking activity (MUA; normalized to ongoing standard deviation across channels and timepoints, within each recording). P-values represent two-tailed (C) or one-tailed (F) t-tests.

Our analyses of LFP oscillatory power used a similar peak centered approach, but was more focused in order to reduce statistical comparisons and to hone in on time/frequency bins with substantial stimulus induced activity (Hamm et al. 2012). We examined the power spectra averaged over all contexts and layers to select time-frequency “regions of interest” (ROIs) of substantial induced activity in a context- and layer-unbiased manner (figure 3A). Six regions of interest were identified (see results). We then assessed DD (deviant vs control) and SSA (redundant vs control) within each ROI, in each of the 4 layers. Unlike CSD and MUA, we observed noticeable induced power across all layers for at least one condition for each of these 6 ROIs. Therefore, we carried out repeated measures ANOVAs with LAYER (1-4) and CONTEXT (control, deviant, redundant) for each time-frequency ROI, except for in the one case where the literature presented a clear hypothesis: deviant stimuli should induce strong superficial theta power. For this, we carried out a separate paired-samples t-test to directly test this prediction.

**Figure 3.**
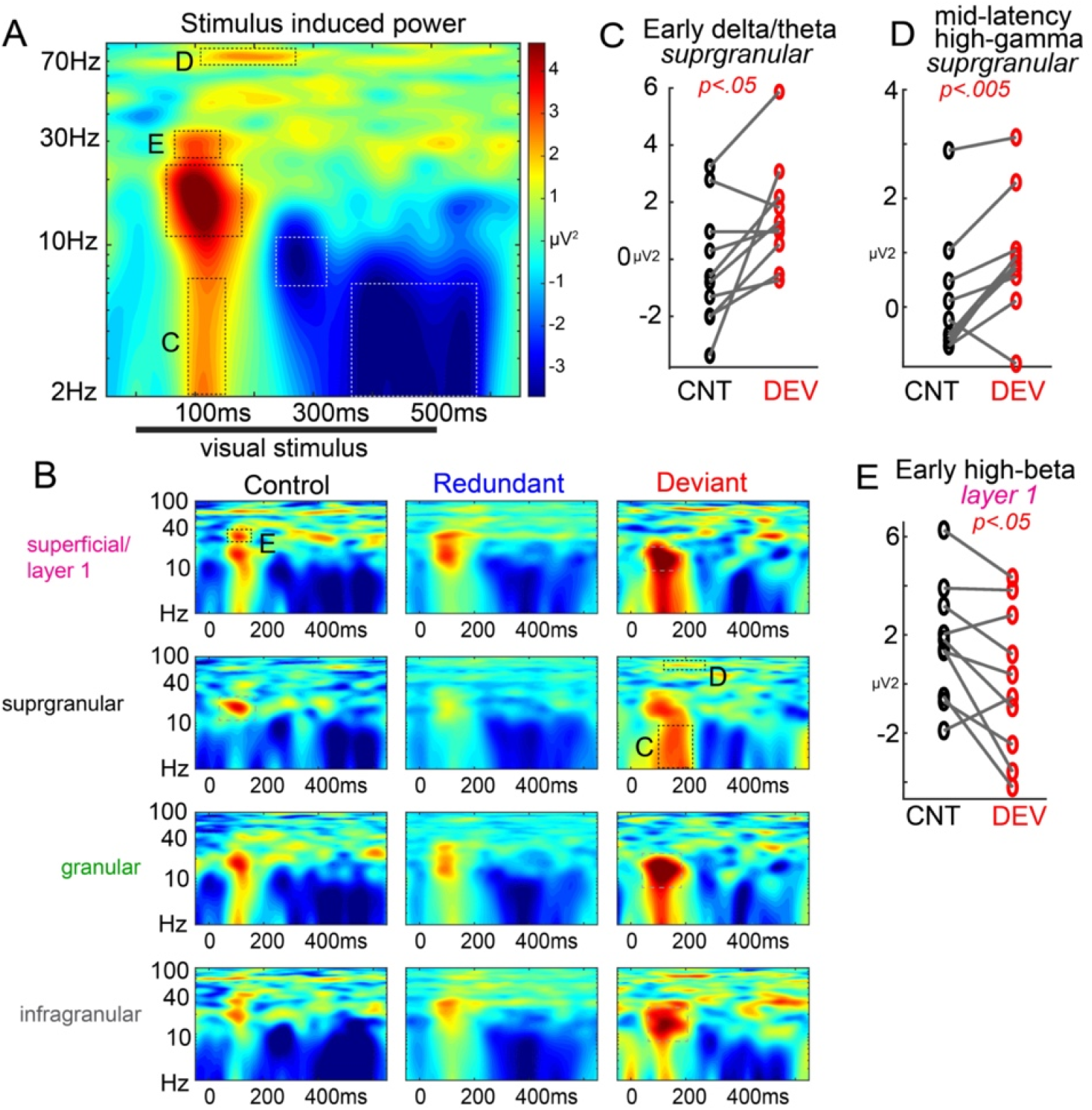
Deviant stimuli evoke a distinct neuro-oscillatory response across neocortical laminae. A) Time-frequency spectral power averaged across all mice, layers, and conditions. Peak/valley regions of interest were identified based on this global average. B) Spectral power averaged for each layer and context condition. C) Early delta/theta power (2-7 Hz) and D) mid-latency (110-260ms) high gamma power (67-76 Hz) was enhanced to deviants in supragranular layers (ROIs indicated in A and B). E) High-beta power (26-36 Hz) was reduced to deviants in superficial/layer 1.

Analyses of IES spectra were more exploratory, as relevant frequency bands, laminar distributions, and interlaminar interactions have not previously been studied in detail at the level of a visual cortical column during the oddball paradigm. We therefore carried out a non-parametric cluster-based permutation framework (Bullmore et al. 1996). One benefit of this method is that it allows empirical generation of differences in spectral content expected under a null hypothesis, achieved by drawing each spectra’s point values from the difference scores upon shuffling condition-labels. If stimulus context does not alter the time-space-frequency landscape of V1 oscillations, we expect the shuffling procedure to approximate this distribution.

Cluster test-statistics were computed based on a ‘cluster-mass’ threshold (Bullmore et al. 1999), selecting for contiguous regions across cortical depth-frequency with context differences (SSA or DD) that significantly differed from clusters arising by chance (i.e. > z_crit_ on the z-scored null distribution). To impose a stringent correction for multiple comparisons, we created a distribution of the maximum cluster-masses from each of the shuffled spectra, and considered clusters in the observed difference maps significant if they were greater than 95% of max null cluster-masses (p<.025 for two-tailed, 2,500 permutations).

### Locomotion Detection

Locomotion, along with measurements of pupil diameter, blinks, and whisker pad movement, was captured at 30 fps during each experiment via a Logitech C920 HD Pro webcam mounted ≈20 cm away from the mouse’s face, illuminated by a dim 617hm LED. These motor variables were extracted and calculated post-hoc by singular value decomposition of manually selected ROIs using the open-source Facemap software (Stringer et al. 2019). Motor traces were analyzed similarly to LFP data, as stimulus-triggered averages C, R, and D (figure S1).

### Histology

Following experiments, mice were deeply anesthetized and transcardially perfused. Extracted brains were cryosectioned coronally for 3D reconstructions of LFP multielectrode tracks using SHARP-Track (Shamash et al. 2018) for registration to the Allen CCF.

## Results

Awake head-fixed mice (figure 1A) viewed sequences of full-field moving square-wave gratings in two separate runs comprised of three visual sequences total: a many-standards control (8 orientations, random order, all p=.125), an oddball (2 orthogonal orientations, with one p=.875) and an oddball flip-flop (figure 1B). This allowed for the analysis of neural responses to the same stimulus when it was contextually redundant, deviant, and rare but not deviant (control; (Harms et al. 2016; Ross and Hamm 2020)). Locomotion, whisking, and blinks did not differ between stimulus contexts (figure S1A-C). Deviant stimuli induced a transient late emerging decrease in pupil diameter emerging approximately 250ms post-stimulus (figure S1D,E). This has not been previously described in the oddball paradigm in rodents, and may serve as a peripheral readout in future investigating causal roles in prediction-error perception.

### Signatures of predictive processing in multiunit activity and current source density across the cortical column

First we focused on how context modulates neural activity in V1 with regard to extracellular currents and population spiking. Augmented neural responses to the stimulus in the “deviant” context (relative to control), were termed “deviance detection” (DD), while reduced responses to the stimulus in the redundant context are termed “stimulus-specific adaptation” (SSA). Our past work with two-photon calcium imaging has showed that at the level of individual neuron somatic outputs, DD is mostly restricted to layer 2/3 in V1 (Hamm et al. 2021), similar to visuomotor mismatch (Jordan and Keller 2020), while SSA is present in most excitatory cells throughout layers, including in layer 4. Our analysis of CSD here showed that early currents in putative granular layers are indeed reduced to redundant stimuli, showing SSA (Fig 2A-C; *t*(9)=-2.41, *p<.05*). Interestingly, this reduction is also present to deviants as well (Fig 2B; *t*(9)=-3.58, *p<.01*). This suggests that the adaptation of thalamocortical inputs in the visual oddball paradigm affects is broad and not stimulus specific. This is suggestive of a type of general adaptation during the oddball paradigm as previously described in V1 (Hamm et al. 2021), which could, in theory, reflect synaptic depression of thalamocortical synapses during the rapid presentation of similar stimuli in the oddball paradigm. We also found a CSD signature of DD slightly later in superficial layers, as expected (Javitt et al. 1996), yet this was only statistically significant at a trend-level (*t*(9)=-1.98, *p=.079)* and was a decrease in overall current (Fig 2B,C). One possibility is that this reflects a somewhat later mismatch between the top-down projections (which synapse superficially in V1) with the bottom-up stimulus information, and thus a transient asynchrony.

We then focused on multiunit activity (MUA), which indexes locally (< 50µm) aggregated neural spiking (figure 2D). As expected, we identified SSA to redundant stimuli in the early time bin (50-80ms) in putative granular layers (figure 2E,F; *t*(8)=-2.02, *p<.05*), consistent with our past findings of SSA in layer 4. The fact that this early granular layer MUA response is not reduced to the deviant stimulus (like we saw a reduction in the CSD) may suggest that while thalamocortical inputs are generally adapted or depressed, the layer 4 neurons which are selective for deviant-orientation correlated on/off fields and/or orientations are not adapted. Thus, the MUA displays a true SSA, while the CSD shows a more general effect. Next, we also found SSA in slightly later superficial/layer 1 (figure 2E,F; *t*(8)=-2.46, *p<.05*) and deeper layer 5 responses (figure S2; *t*(8)=-2.09, *p<.05*) in the next time bin (80-110ms), suggesting these may be the next populations innervated by adapted layer 4 neurons.

Following this, we identified DD responses in putative layer 2 neurons at approximately 140-170ms (*t*(8)=2.44, *p<.05*) and then slightly later in deeper, putative layer 3 from 170-230ms (*t*(8)=5.18, *p<.001*). This is consistent with our past work and, interestingly, shows that DD occurs later than SSA and in spatially distinct neural populations. Interestingly, the average MUA responses suggest a deeper signature of DD in putative layer 5a (figure 2E) concurrent with the layer 2 DD response, but upon further examination, this difference in means is strongly driven by only 2 recordings and does not reach statistical significance at the population level (figure S2; *t*(8)=1.54, *p=.08)*. One possibility is that those MUA recordings in particular captured largely inhibitory neurons in layer 5, which are known to be innervated by layer 2/3 excitatory cells (which potentially exhibited DD). Future work looking at how different neural populations exhibit DD and SSA is warranted.

In general, CSD activity did not show large effect sizes or widespread differences for DD or SSA as compared with MUA. This may stem from that fact that CSD is inherently more variable, with lower signal to noise ratio, and requires more than the ≈10 trials per condition we employed to generate stable waveforms. In this study, fewer trials were used to balance longer term adaptation in the paradigm. Our smaller CSD may also stem from the fact that moving grating stimuli were employed, which could ensure more local neural firing but lead to cancelling phase offsets that distort CSD when averaged as a waveform. Analysis in the time-frequency domain affords a solution to this issue, and also has shown more reliable effects in clinical neurophysiology studies (Javitt et al. 2020).

### Deviant stimuli induce a distinct neuro-oscillatory signature in supragranular V1

Next, we turned to analysis in the time-frequency domain. Narrowband brain rhythms supporting cognitive and behavioral functions (e.g. theta, gamma) tend to be preserved in their spectral properties despite astronomical differences across mammalian taxa (Buzsáki et al. 2013). We sought to test whether or not contextually-evoked response differences correspond to specific neuro-oscillatory bandwidths in a predictive coding framework. Rhythm-based models of predictive coding are a distinct class which posit a neurophysiological implementation for the bi-directional flow of cortical predictions and prediction-errors, involving distinct oscillatory channels (Arnal and Giraud 2012; Bastos et al. 2012, 2020). These models divert away from specialized prediction-error circuits, favoring instead the notion that common, ‘canonical’ cortical circuitry used for many functions can be gated by ongoing oscillations that facilitate or inhibit the processing of cortical input. Predictions about upcoming signals originate in deep layers of hierarchically higher cortical regions and terminate in layer 1 of sensory regions and occupy alpha-beta frequencies, while prediction errors originate in layer 2/3 and are fed-forward to update higher brain areas via gamma (Lundqvist et al. 2016; Bastos et al. 2018) and theta (Bastos et al. 2015) oscillations. We asked whether predictive processing under a visual oddball paradigm would elicit the same basic patterns in V1.

To simplify analyses and avoid biases, we focused on time-frequency “regions of interest” (ROIs) in the grand average power spectra (averaged across layers and conditions; Figure 3A). These ROIs were early delta/theta (2-7 Hz; 100-150ms), early low beta (12-22 Hz; 65-180ms), early high beta (26:35 Hz; 90-120ms), mid-latency high gamma (68-77 Hz; 110-260ms), mid-latency alpha desynchronization (6-12 Hz; 240-310ms), and late delta/theta desynchronization (2-7 Hz; 350-560ms). We then assessed DD and SSA within each of the 4 layers.

Past work (primarily in the auditory domain) has shown that the deviance detection signals in humans and rodents that DD is associated with increases in stimulus induced/evoked low theta power (2-8Hz; (Javitt et al. 2018; Lee et al. 2018)), and on our MUA results (and other work) suggests that layer 2/3 shows strong DD (figure 2F). Thus, we first examined the delta/theta ROI. As expected, layer 2/3 exhibited enhanced theta power to deviants (*t*(9)=2.84, p<.05; figure 3C), but not redundants (t(9)=0.46, p=.656). This was not seen in any other layer (all p>.44).

Examining these time-frequency ROIs more broadly, we also identified a layer by context interaction effect for high mid-latency gamma oscillations (f(2,3)=2.29, p<.05; figure 3D) which was driven by increases specifically in layer 2/3 to deviant stimuli (*t*(9)=4.53, p<.005; all other layers and comparisons p>.24). This effect corresponded in time and space to the late MUA DD signal (figure 2F). We also identified a layer by context interaction effect for early beta oscillations (f(2,3)=3.06, p<.01; figure 3E) which was driven by decreases specifically in layer 1 to deviant stimuli (*t*(9)=-2.58, *p*<.05; all other layers and comparisons p>.29). This effect corresponded in time and space to our deviant related decrease in superficial current source (figure 2B).

No other statistically significant effects of stimulus context or layer by context interactions were identified for other time-frequency ROIs. However, examining the average power time-frequency spectra (figure 3B), the early beta response, present strongly across all layers to the onset of stimuli, appeared strongest to the deviant in layers 1, 4, and 5, but was weaker to deviants in layer 2/3. This pattern was present in all mice except for one (figure S3). Carrying out the layer by context ANOVA without this mouse included revealed a trend-level stimulus by context interaction (*F*(2,3)=1.92, *p*=.095), driven by deviant stimulus increases in this low-beta power in layers 4 and 5 (*p*<.05). This mouse was curiously not an outlier in any other measure of interest, and exclusion did not substantially impact any other effect in the study. Further, examining scatterplots for all other time-frequency ROI, layer, and condition did not reveal any other such potential outlier-driven effects or non-effects. This effect, though consistent with beta oscillations originating in feed-back circuits, should be interpreted with caution.

In sum, processing of contextually deviant stimuli in the oddball paradigm involved increases in theta- and gamma-band oscillations in layer 2/3. These frequency bands and this layer of cortex are believed to carryout feed-forward processing, which is consistent with deviance detection reflecting a “prediction error” which is fed-forward in cortical circuits. Further, deviant stimuli also involved a decrease in high beta-band power in layer 1. Beta oscillations are believed to be the preferred frequency bandwidth for cortical feedback, and feedback cortical projections terminate in layer 1. Thus, this temporary disruption of layer 1 beta is consistent with deviant stimuli mismatching contextual “predictions”.

### Deviant stimuli desynchronize low-gamma coherence between layer 1 and layer 2/3

Lastly, we sought to examine how deviant stimuli alter local synchrony across the cortical columnar circuit by looking specifically at inter-laminar phase coherence. Synchrony in neural circuits serves a diverse set of functions from temporal binding to dynamic grouping of features for joint processing (Singer 1999). We focused our analysis on inter-laminar synchrony between superficial-layers (i.e. layer 1) and all other electrode contacts to study. We were mainly interested in how local processing across lamina in V1 related to ongoing “predictive” information, which theoretically arrives in layer 1 (Douglas and Martin 2004). In essence, synchrony to the “top-down” component. We estimated inter-laminar synchrony using the mean phase consistency at each electrode when contacts 1 and 2 were used as reference seeds. Spectra for each stimulus context showed strong lay1-lay2 and lay1-lay5a synchrony in the theta/alpha band for the first 250ms the stimulus was on-screen (Figure 4A). We conducted two-tailed cluster-based permutation testing on deviant- and control-evoked synchrony differences, given we had no *a priori* hypotheses about the direction in which stimulus context might exert its effect. This analysis revealed a significant low gamma-band desynchronization (34-41 Hz) to deviant stimuli in ventral layer 3/dorsal layer 4 relative to superficial layer 1 (Figure 4B; p<.025). Closer inspection of this trend for each individual mouse showed this decrease in synchrony was robust in 9 out of 10 mice. We also examined the “bottom-up” component, applying the same procedure when using the granular layer (contacts 8 and 9) as the reference seed. There were no robust inter-laminar synchrony differences from this perspective.

**Figure 4.**
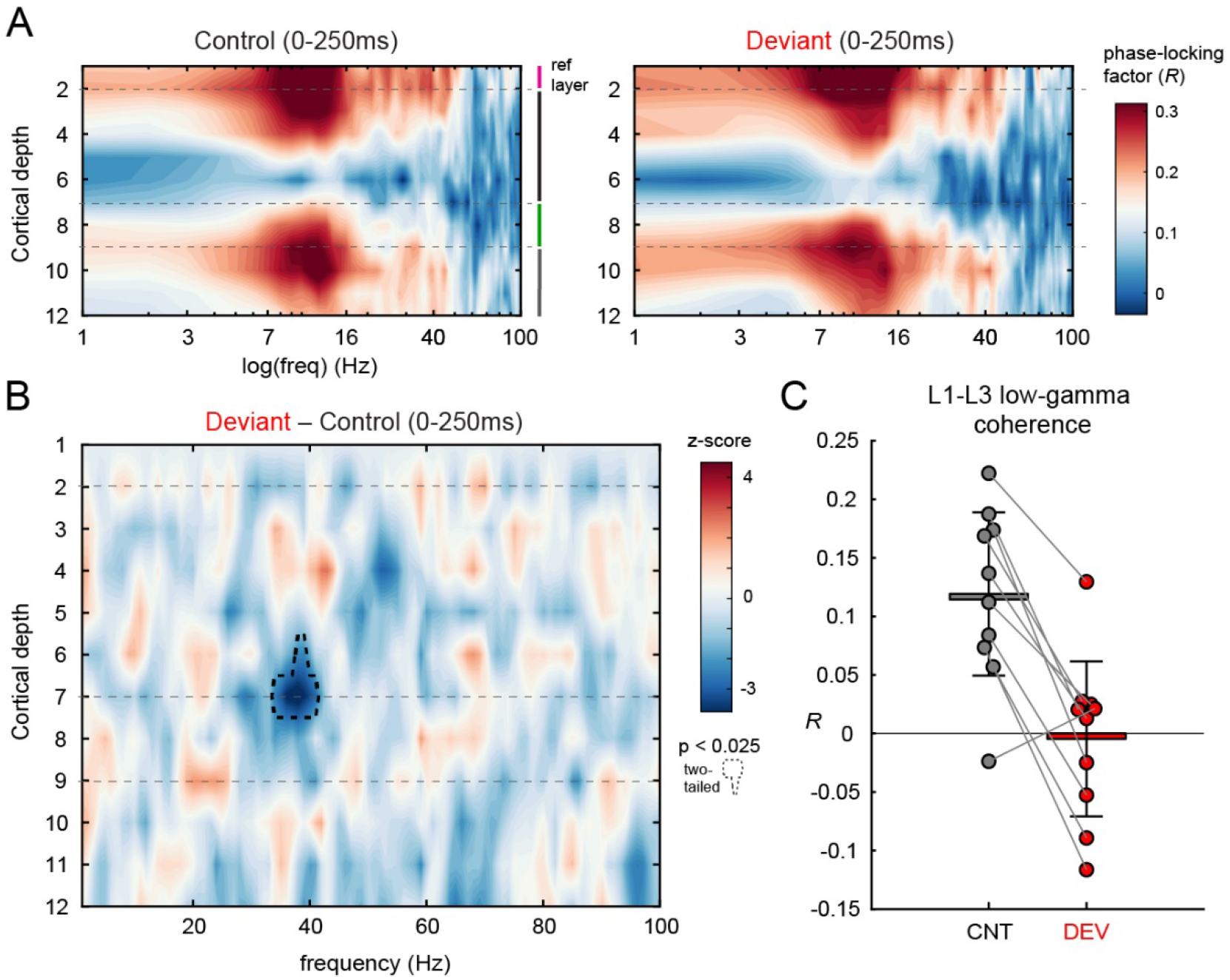
Deep layer 3 desynchronizes with superficial layers in deviant contexts. A) Inter-laminar synchrony maps for the control (left) and deviant (right) stimulus contexts, referenced to superficial cortical layer 1 (average across electrodes 1 and 2), and plotted on the log-scale to emphasize theta-alpha synchronization across supra- and infragranular layers. B) Represents the difference map, constructed by subtracting control stimulus responses from responses to the deviant. Cluster-based permutation testing revealed two-tailed significance of the contoured cluster (dotted black line; 34-41 Hz). C) Depicts the average phase-locking factor (*R*) of points in the contoured region (colored line) for each of the 10 mice, where black bars represent the standard deviation.

## Discussion

We studied V1 LFPs from 10 mice presented with full-field visual gratings in various sequences, aiming to characterize the spatiotemporal dynamics associated with deviance detection (DD) with respect to cortical laminae, frequency bands, spiking, and global transsynaptic currents. We further examined whether a predictive coding framework accurately describes cortical activity evoked during a classic visual oddball paradigm. Our results generally concord with this framework, and enhance basic understanding of how cortical circuits respond to contextually deviant stimuli – an important piece of information for interpreting a large clinical literature on “mismatch negativity” (MMN), an EEG analogue of DD measurable in humans.

Results suggest that unpredicted stimuli – i.e. “deviants” in an oddball paradigm – evoked robust responses in supragranular layer 2/3 (L2/3), arising after 100ms post-stimulus. While early work on translating human MMN suggested that rodents may only exhibit stimulus-specific adaptation (SSA; i.e. simple reduction in responses to repeated stimuli without a separate MMN-like augmented response to deviant stimuli), rigorous studies employing additional paradigms (e.g. the many-standards control) have since conclusively demonstrated that rodents exhibit both auditory and visual DD. Our work further solidifies this case and clarifies that SSA is spatially, temporally, and neuro-oscillatorily distinct from DD. Signatures of SSA arise early (≈50ms) in granular L4 CSD and MUA, while signatures of DD first involve delta and beta oscillatory disruptions in L2/3 and L1, followed by MUA activity in L2/3. This suggests that SSA may be present either in reduced thalamocortical terminals (synaptic depression) or in some adaptation in post-synaptic integration in post-synaptic L4 granule cells. In either case, SSA is likely mechanistically distinct from DD, but whether DD is functionally dependent on SSA remains an open question. That is, for DD to occur, it is not known whether a locally adapted neural population need to be present for, e.g., lateral disinhibition to indirectly “prime” a separate population of cells which carry out DD (Ross and Hamm 2020). Future work should investigate this possibility with more complex paradigms and single cell stimulation/inhibition.

DD signals included early increases in low frequency oscillations (in the traditional “delta/theta” range; 2-7 Hz) and later (>100ms) spiking and high gamma-band power. This laminar and time-frequency signature is consistent with DD representing “prediction errors” in feed-forward circuits. L2/3 is commonly the source of “feed-forward” projections to hierarchically “higher” cortical regions (Douglas and Martin 2004). Theta and gamma oscillations have also been recently shown to coincide with feed-forward signaling in hierarchical cortical networks (i.e. from lower brain regions to higher brain regions (Bastos et al. 2015, 2020)). Notably, prediction-errors are thought to be primarily “fed-forward” in cortical networks to update internal models of the environment maintained in higher cortical regions (Bastos et al. 2012).

On the other hand, beta oscillations have been shown to organize top-down or “feed-back” signaling in cortical networks (Bastos et al. 2015). In the current study, we found decreases in early stimulus-induced “high beta” in layer 1 to deviant stimuli. One interpretation of this is that sensory data about the current stimulus and the predicted stimulus are integrated in layer 1, where dendrites of V1 pyramidal cells receive top-down inputs from higher cortical regions. The mismatch of the deviant sensory data (from the bottom-up/feed-forward direction) with the predicted sensory data (from the top-down/feed-back direction) may have temporarily disrupted this “predictive” high-beta oscillation in layer 1. Thus, this beta disruption could be an early signature of deviance detection (occurring prior to MUA firing and gamma-oscillations in L2/3) that signifies the momentary disruption of predictive processing in the visual system, leading to subsequent firing and gamma-band activity in L2/3. Future work could more directly test this model by suppressing top-down inputs to V1 with optogenetics at this particular timepoint/frequency band to see how subsequent DD signals are affected.

Interestingly a spatiotemporally and spectrally distinct “low-beta” response was present as well (figure 3B). This did not statistically differentiate conditions, but many mice showed stronger low-beta responses to the deviant stimulus in deeper layers (figure S3). While this spatio-spectral distribution is consistent with feed-forward gamma/theta and feed-back beta discussed above, it is curious that such a DD signal is present in deep layers: a point which is not entirely consistent with a simplified microcircuit model of predictive processing (as deep layers send cortical feed-back, and superficial layers send cortical feed-forward signals). Curiously, there was not a concomitant multiunit firing response with this DD signal in deep layers (figure 2) and our past work with two-photon calcium imaging also shows that the majority of layer 5 neurons do not show DD (Hamm et al. 2021). One possibility is that deviant stimuli induce an inhibition of layer 5 feedback projections, which shows up in the low-beta band (a putative preferred frequency band for SST+ and VIP+ interneurons; (Veit et al. 2017; Van Derveer et al. 2020; Bastos et al. 2023)) and which showed up in a small subset of our mice in the MUA signal (figure S2). Interneurons are more spatially sparse in the cortex, a point which could explain the variability across mice (as our electrodes might have differentially sampled one spatially proximal population vs another by chance). Future work with optotagging or deep-calcium imaging of specific interneurons should investigate this possibility.

A rich foundation of literature supports the interplay between gamma and theta as a key motif of interlaminar and inter-areal neuronal communication (Fries 2005, 2015; McGinn and Valiante 2014). Our results are consistent; we observed strong L2 and L4/5 theta-alpha resonance across all stimulus contexts in this paradigm. Further, we identified a decrease in low gamma coherence between layer 1 and deep layer 3 that spatiotemporally aligned with where we saw deviant-related changes in supragranular/granular MUA. In thinking about how this maps onto the microcircuit architecture of V1, one possibility is that it could reflect a mismatch between predictions (arriving in layer 1) and feed-forward sensory data (arriving in basal dendrites of pyramidal cells in deep layer 3) which is unique to the deviant stimulus condition. Another possibility, given the frequency-characteristics of this effect, is that deviant stimuli disrupt PV+ interneurons in the early phase of presentation, which have been shown to support feed-forward sensory processing and have been well-characterized as gamma generators (Cardin et al. 2009).

Our designation of “low-theta/delta” band as 2-7Hz was driven by our evoked responses (figure 3A). Intriguingly, the theta-band has been found to be slower (1-4Hz) and more transient in humans (Jacobs 2014; Burke and Maurer 2020; Foo and Bohbot 2020), compared with sustained activities ≥8Hz in rodents (Vanderwolf 1969; Watrous et al. 2013). On the other hand, some work in mouse V1 suggests that alpha-like rhythms may occupy frequencies in the 3-6Hz range (Senzai et al. 2019). Nevertheless, differences in precise peak-frequency may not detract from similarity of function. Indeed, these oscillations still exhibit congruent dynamics supporting the same cognitive activities despite their frequency-shift across species (e.g. spatial navigation (Watrous et al. 2013) and REM sleep (Jacobs 2014)), as well as recruit other known key players (e.g. gamma) for cross-frequency coupling interactions (Lisman and Buzsaki 2008; Clemens et al. 2009; Ferrara et al. 2012). Such areas of convergence support the potential generality of this neuro-oscillatory marker in MMN-generation across mammals.

Interestingly, while we identified CSD signatures of SSA and DD in L4 and L1, respectively, we did not find large CSD signatures of DD in layer 2/3 or other layers, however, which is somewhat inconsistent with our past study mouse V1 DD (Hamm and Yuste 2016). One possible explanation is that CSD is inherently more variable (especially with lower trial numbers) and, importantly, it may be more sensitive to stimulus parameters. In our past work we employed static gratings while we used moving gratings here. A difference in how these stimuli activate different regions of retinotopic V1 as they move (or do not move) across the visual field may give rise to variability in the CSD which is difficult to model or expect. However, moving gratings may more consistently evoke spiking (by avoiding gaps in the visual field), and analysing time-frequency power and phase-locking effectively circumvents these problems by dissociating measures of “stimulus-evoked” (or locked) from “stimulus-induced” dynamics.

Given the instability of superficial cortical multielectrode probe recordings in awake moving animals, collecting data on single units (spiking from individual neurons) was challenging. Our goal here was to match, replicate, and extend the understanding of how different lamina adapt and detect deviance in V1, and also to uncover the neurooscillatory landscape within and between cortical layers first, especially insofar as findings in this laminar/CSD domain can still relate to EEG and LFP recordings in humans and non-human primates. Further, novel insights into disease relevant circuit dysfunction could be gleaned by studying the intracortical dynamics identified here in one of the variety of mouse lines which recapitulate genetic or cellular aberrations commonly found in schizophrenia and other diseases known to involve altered MMN (Featherstone et al. 2015; Hamm et al. 2017, 2020).

## Acknowledgments

We would like to thank the staff of GSU’s Division of Animal Resources for their consistent support in animal care and colony management, as well as Lab Manager Antanovia Ferrell for the upkeep and maintenance of lab equipment that makes this work possible. This work was funded by the National Eye Institute (R01EY033950), National Institute of Mental Health (K99/R00MH115082), Brain and Behavior Research Foundation (YI30149), and the Whitehall foundation (2019-05-443). All materials are commercially available. The authors declare no competing financial interests.

Correspondence should be directed to Jordan Hamm, 813 Petit Science Center, 100 Piedmont Ave, Atlanta, GA 30303; 404-413-5398;

## Author contributions

JPH conceptualized the experiments. CGG performed the electrophysiological and behavioral recordings, histology, and microscopy. DR and CGG analyzed behavioral data. CGG and JPH analyzed electrophysiological data, made the figures, and wrote the manuscript.

## Data and code availability

Raw data has been made publicly available as a part of the Open Science Framework (Gallimore and Hamm 2022). Custom processing and analysis scripts necessary to replicate these results will be uploaded to GitHub at the time of publication.

