## Supplemental figures and methods for "Spatiotemporal dynamics across visual cortical laminae support a predictive coding framework for interpreting mismatch responses"

**Supplemental Info**

Supplemental Methods

Non-parametric cluster-based permutation testing provides an elegant way of circumventing the multiple comparisons problem (Maris and Oostenveld 2007) when handling high-dimensional datasets that contain meaningful autocorrelative temporal structure. Our choice of 2,500 (t-f power, IES) shuffles for cluster-based permutation testing was justified by a prior simulation study by Pernet et al. (2015), where the authors concluded stability of results is achieved ≈800 permutations with diminishing returns beyond (Pernet et al. 2015). Using the maximum cluster-level statistic to calculate p-values under the permutation distribution is essential to control the false alarm rate for all clusters, despite its reduction in sensitivity for smaller clusters (Maris and Oostenveld 2007). This expense is actually preferred here, as cluster-mass (c-m) distributions were log-normal in all cases (Fig S5, S8), consistent with other metrics of biological size (Huxley 1932) and functional physiological quantities (Mizuseki and Buzsáki 2013; Buzsáki and Mizuseki 2014; Scheler 2017). Consequently, the smallest identified clusters in observed difference maps (prior to correction) corresponded to exceedingly small, transient regions of activation in time-frequency (t-f) space, and were most likely spurious positives, whereas the largest identified clusters corresponded to sustained activations in t-f space representing the contextually-evoked neuronal signals of interest. This statistical approach is further merited by the c-m significance threshold's basis in a null distribution derived empirically (through shuffling), allowing the method to flexibly scale with t-f decomposition parameters chosen *prior* to statistical analysis (e.g. wavelet width) and obviating any arbitrary experimenter definition of t-f bins (Cohen 2021).

### Supplemental figures

Figure S1

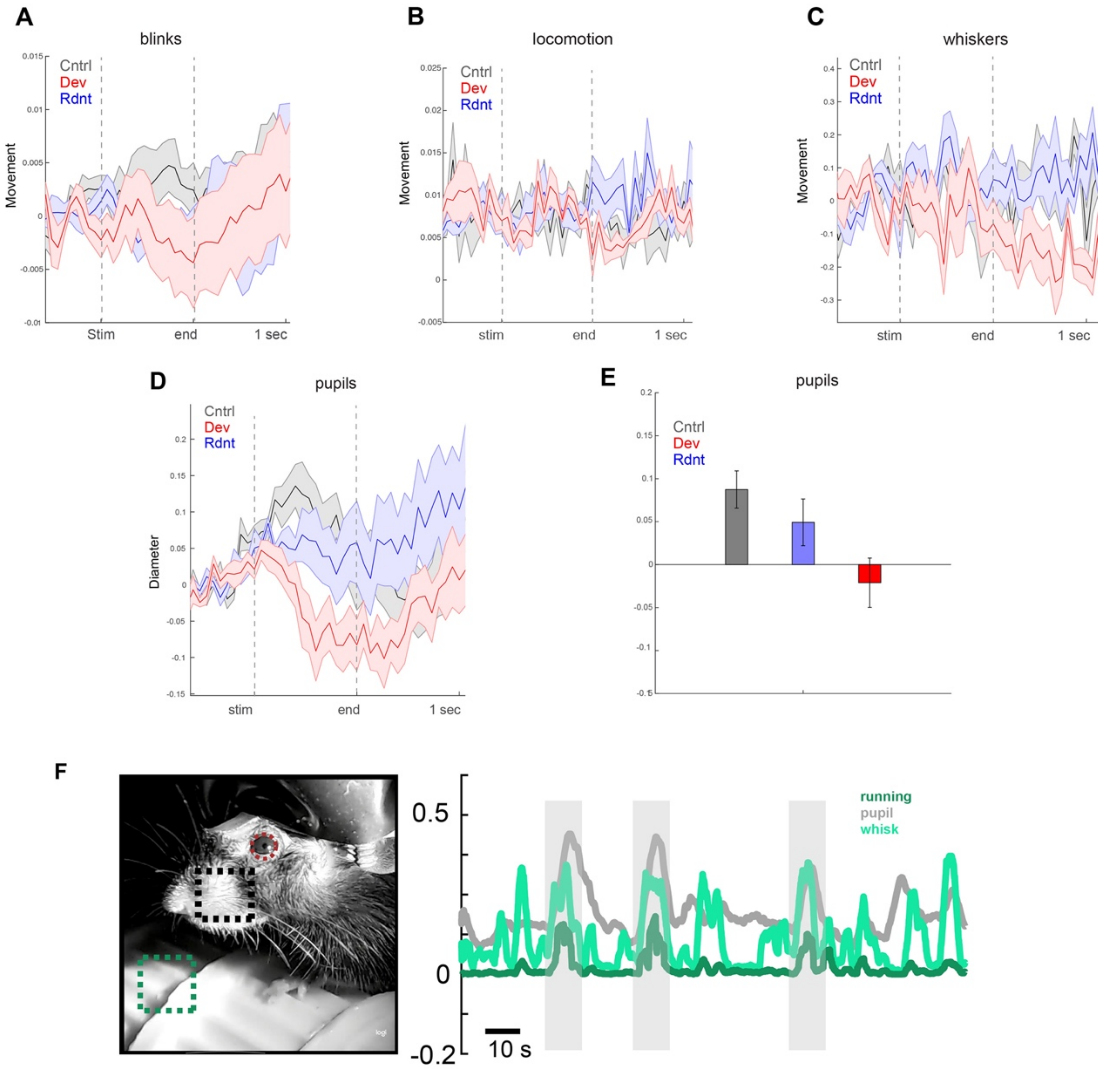


**Figure S1.** a) blinks (indicated by change in total motion energy of eye region) averaged across mice and trials, from 0.5 second before to 1 second after stimulus onset. No effect of stimulus context was observed for blinks (repeated measures ANOVA; $F(5)=0.95$, $p=0.42$). b) locomotion (indicated by change in total motion energy of running wheel; repeated measures ANOVA; $F(5)=0.01$, $p=.99$), or c) whisker pad movement (indicated by change in total motion energy of whisker pad; repeated measures ANOVA; $F(5)=0.55$, $p=0.59$). d) Pupil diameter (indicated by change in size of facemap pupil ROI) on the other hand showed a significant effect of context during the stimulus interval (panel e, repeated measures ANOVA; $F(5)=4.2$, $p<.05$). f) Example frame of mouse face video, displaying behavior variable ROI's for movement analysis (left). Change in pupil, running, and whisking over time course is shown (right). All error bars denote SEM (a-d).

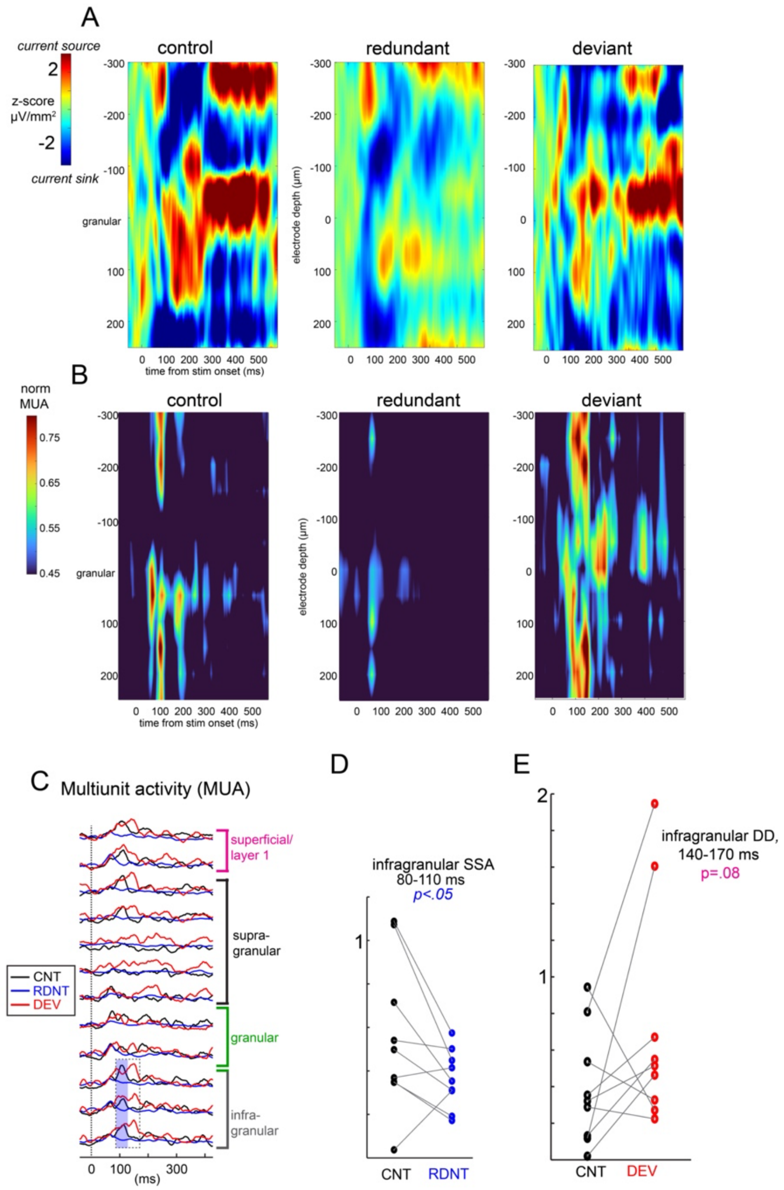


**Figure S2. Supplementary CSD and MUA plots.** A) trial and mouse averaged CSD for each condition. Notably, this diverges from the average rectified (AVREC) montage shown in figure 2. This is because averaging CSD over mice and trials without first rectifying of the currents (taking the absolute value; i.e. converting to AVREC) leads to cancelling if when sinks and sources do not match up across mice and trials. This affects the redundant condition more, as there are more redundant trials. We present this here mainly to demonstrate that the distributions of sinks and sources are generally the same between conditions.  B) Trial and mouse averaged MUA and for each condition. C) same data as in B, but represented in line form. D) Statistically significant infragranular SSA, occurring later than layer 4 SSA. E) large MUA peaks to deviant in infragranular layers driven by two mice.

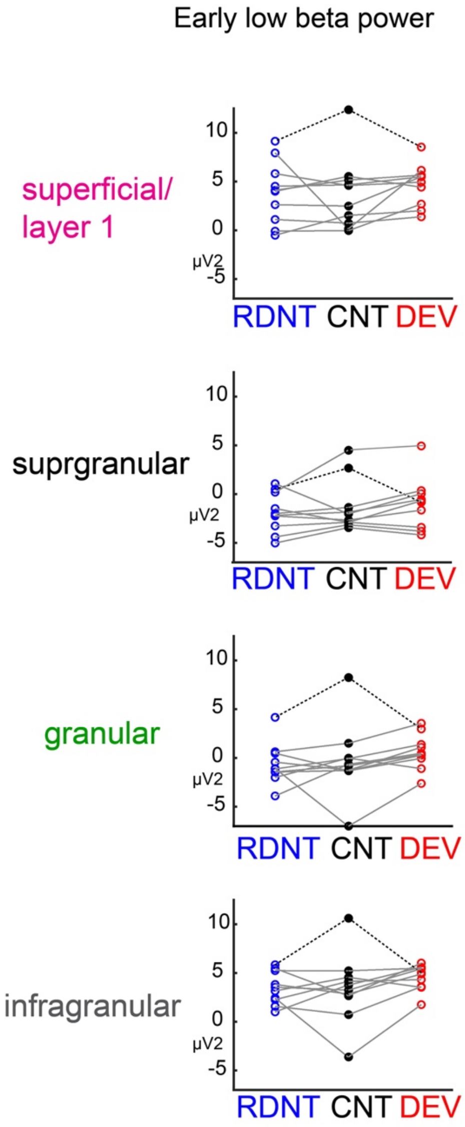


**Figure S3. Scatter plots for early low beta power.** Each point represents a mouse's averaged induced power at 12 to 22 Hz, from 65-180ms post-stimulus onset. This was the largest amplitude response across all layers/mice. There was no significant context main effect of context by layer interaction, but a single outlier mouse (dotted lines above), with strong responses to the control condition, seemed to influence these statistics. Without this mouse, there was a trend-level layer by context interaction ($p=.07$), with significantly/trend-level greater power in granular ($t(8)=3.34$, $p<.05$), infragranular layers ($t(8)=2.97$, $p<.05$), and superficial/layer1 ($t(8)=2.12$, $p=.06$) to deviant stimuli.
